# Identification and characterization of a novel exo-(N-)glycanase activity of NGLY1 on ENGase-digested N-GlcNAc proteins *in vitro*

**DOI:** 10.1101/2025.08.21.671495

**Authors:** Yilin Ye, Xinrong Lu, Shaoxian Lyu, Yuxin Zhang, Qiange Lin, Xin Qian, Junyou Lin, Zhijian Yu, Li Chen, Guiqin Sun

## Abstract

N-Glycanase 1 (NGLY1) is known for its activity to completely remove conjugated N-glycans from glycoproteins. Early studies illustrated that NGLY1 defects were associated with a rare disease named NGLY1-related congenital disorder of deglycosylation (NGLY1-CDDG). Although extensive research has been conducted over the past decade on the biological impact of NGLY1’s endo-(N-)glycanase activity in cells and on disease, whether NGLY1 also exhibits an exo-(N-)glycanase activity remains open. In this study, an exo-(N-)glycanase activity of NGLY1 on the N-GlcNAc proteins was firstly reported and characterized *in vitro*. To distinguish for the conventional endo-(N-)glycanase activity and the newly reported exo-(N-)glycanase activity of NGLY1, the active sites of exo-NGLY1 were predicted and compared with those of endo-NGLY1. In addition, the correlation between NGLY1’s exo-(N-)glycanase activity *in vitro* and NGLY1-CDDG’s clinical characterization was investigated and discussed. The observation of NGLY1’s novel exo-(N-)glycanase activity expanded the functional repertoire of NGLY1 beyond canonical endo-(N-)glycanase-mediated deglycosylation, and may provide a new framework to explain the clinical heterogeneity of NGLY1-CDDG.

## Introduction

N-Glycanase 1 (NGLY1) is an enzyme responsible for cleaving oligosaccharide moieties from misfolded glycoproteins to enable their proper degradation(1). It could hydrolyze the amide bond between the proximal N-acetylglucosamine (GlcNAc) residue and the asparagine (Asn) side chain, thereby removing the N-glycans from the glycosylated proteins in the cytoplasm and releasing the complete oligosaccharide chain, converting the asparagine (Asn) on the peptide chain into aspartic acid (Asp)(1–3). In 1993, Suzuki T et al reported the activity of NGLY1 (cytosolic PNGase)(4). It was discovered that NGLY1 participated in the degradation of misfolded glycoproteins during the process of glycoprotein synthesis through the endoplasmic reticulum associated degradation (ERAD) pathway, thereby maintaining normal cell functions(5). In 2013, it was reported that NGLY1 deficiency might be considered the first “congenital disorder of deglycosylation (CDDG)"(6), with the main clinical manifestations including developmental delay, intellectual disability, absence of tears and less sweating, abnormal liver function, and motor dysfunction. In 2017, it was reported that NGLY1 deficiency was described as NGLY1-CDDG, and its clinical phenotypic spectrum was expanded(7).

Early studies suggested that NGLY1 was essential for ERAD, and the compromised *Ngly1* activity for ERAD could be partially complemented by *Engase* mediated deglycosylation of misfolded glycoproteins. Early studies demonstrated that NGLY1 was a component of the endoplasmic reticulum-associated degradation (ERAD) machinery(8,9). This study identified a novel exo-(N-)glycanase activity of NGLY1. We hypothesized that this activity could partially compensate for the deglycosylation of misfolded glycoproteins mediated by ENGase. In 2017, it was reported that homozygous *NGLY1* knockout was embryonic lethal in mice under C57BL/6 background and this phenotype could be rescued by *Engase* knockout(10). To explain the phenotype, it was proposed that elevated ENGase activity may cause excess formation of N-GlcNAc proteins in cell, and the situation was bypassed in *Ngly1* and *Engase* double knockout(11). Yoshida found that deletion of *Engase* partially suppressed the lethality of *Ngly1-KO* and ENGase might be needed to remove bulky N-glycans to allow efficient ubiquitination of Nrf1 by SCF^FBS2^-ARIH1(12). Although studies have shown that NGLY1 deficiency leads to N-GlcNAc proteins aggregation, it is still unclear which enzyme have the exo-(N-)glycanase activity to remove N-GlcNAc proteins.

This article investigated the new exo-(N-)glycanase activity functions of NGLY1. It was discovered that NGLY1 had a new function of removing N-GlcNAc from glycosylated proteins by virtual molecular docking and lectin blot methods, as well as the sites that affected this new function. Additionally, we studied the influence of NGLY1’s exo-(N-)glycanase activity in NGLY1-CDDG.

## Results

### NGLY1 new function——exo-(N-)glycanase activity

In this study, a prokaryotic expression system for NGLY1 was successfully constructed, enabling the purification of the NGLY1 protein from *E.coli* BL21 (DE3). Enzymatic activity assays demonstrated that NGLY1 exhibits endo-(N-)glycanase activity, as evidenced by its action on the high-mannose type standard glycoprotein substrate RNase B which was a conventional substrate of NGLY1 (Fig. 1A). As controls, other N-glycanases were employed, including the prokaryotic-expressed PNGase F and PNGase F-II from *Elizabethkingia meningoseptica* discovered in 2015 by our group(13), both of which also had enzymatic activity toward RNase B (Fig. 1A).

**Figure 1.**
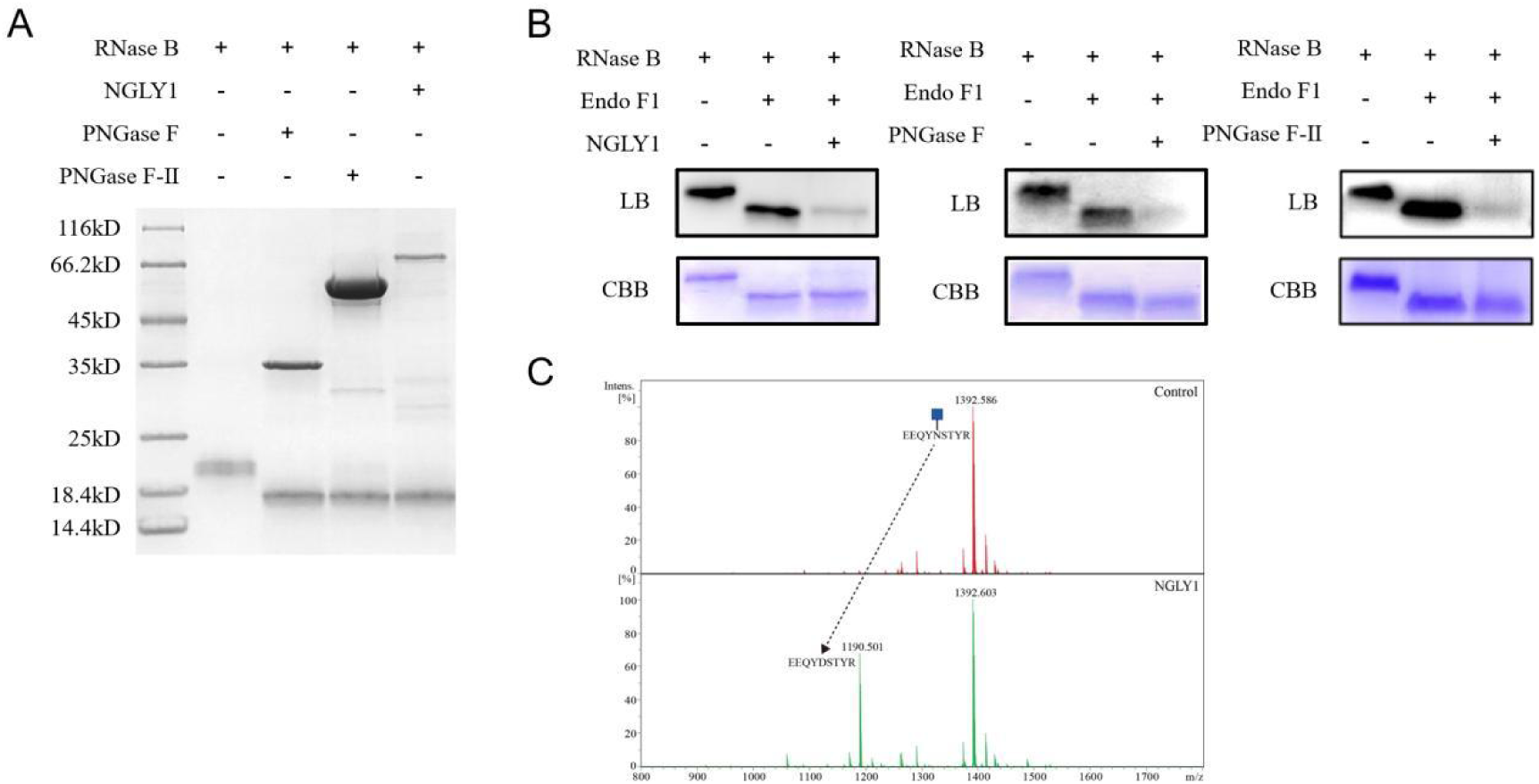
The endo-(N-)glycanase and exo-(N-)glycanase activities of NGLY1. A. N-glycanase’s endo-(N-)glycanase activity. The first lane showed the band of RNase B alone; the second lane showed the bands of RNase B substrate treated with PNGase F, PNGase F-II and NGLY1 respectively. The downward shift of the RNase B band indicated that all three enzymes have deglycosylation functions. B. N-glycanase’s exo-(N-)glycanase activity. The first lane showed the band of RNase B alone; the second lane showed the bands of RNase B substrate treated with Endo F1; the third lane showed the new substrate (enzymatic products of Endo F1 cleaving RNase B) treated with NGLY1, PNGase F and PNGase F-II respectively. C. Mass spectrometry analysis on NGLY1’s exo-(N-)glycanase activity. The glycopeptide EEQYN(GlcNAc)STYR was incubated with NGLY1 and analyzed by MALDI-TOF mass spectrometry. The resulting spectra displayed control and experimental peaks (red and green, respectively), with all peaks annotated by peptide sequence and molecular ions were present in sodiated form ([M + Na]^+^). CBB: Coomassie Brilliant Blue; LB: Lectin Blot. The darker the color of the band in the lectin blot, the higher the content of GlcNAc.

Following treatment of RNase B with Endo F1 (endo-beta-N-acetylglucosaminidase F1) from *Elizabethkingia meningoseptica*, the N-GlcNAc proteins were obtained and then used as substrates for NGLY1. The results showed that NGLY1 could hydrolyze a single N-GlcNAc on protein. At the same time, PNGase F and PNGase F-Ⅱ also had this new function (Fig. 1B).

To further validate NGLY1’s exo-(N-)glycanase activity, we performed mass spectrometry analysis using the glycopeptide substrate EEQYN(GlcNAc)STYR (m/z 1392)(14). Following NGLY1 digestion, a product peak at m/z 1190 was detected, corresponding to the deglycosylated peptide EEQYDSTYR, while residual substrate remained(Fig. 1C). This demonstrated that NGLY1 hydrolyzes single N-GlcNAc residues from glycopeptides, converting the N-GlcNAc-linked asparagine to aspartic acid and thereby confirming its exo-(N-)glycanase activity. These results were consistent with our lectin blot findings.

### Prediction of the amino acid residues affecting NGLY1’s activity

The three-dimensional structure of NGLY1 was predicted using AlphaFold2 (Fig. 2A), which contained three domains: PUB domain, PNG Core domain, and PAW domain. PNG Core domain was the main catalytic domain. The structures of molecular ligands were designed, and the molecular docking results with NGLY1 were visualized by PyMOL(Fig. 2B and 2C). It was found that the binding between the ligand and the protein mainly occurred in the middle region between PAW and PNG Core (the red box in Fig. 2A).

**Figure 2.**
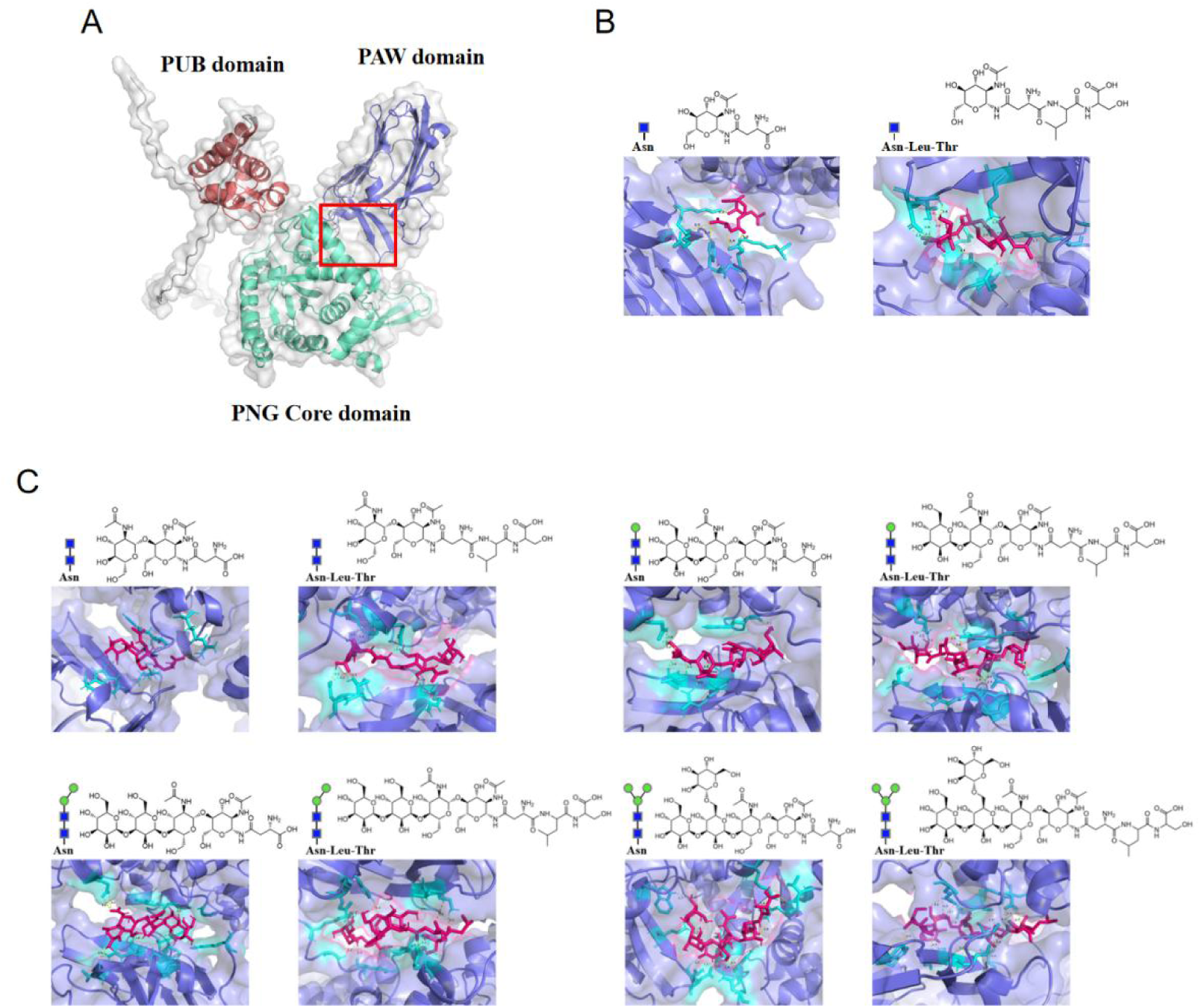
The molecular docking of ligands and NGLY1. A. The 3D structure of NGLY1. The red box was the docking pocket. B. The ligand structure and molecular docking picture to predict the amino acid residues that affected exo-(N-)glycanase activity. Each group of pictures included a ligand schematic diagram, the molecular structure of the ligand, and a molecular docking diagram. In the molecular docking diagram, the bright pink area represented the structure of the ligand. C. The ligand structure and molecular docking picture to predict the amino acid residues affecting NGLY1’s exo-(N-)glycanase activity. Blue square: N-acetylglucosamines (GlcNAc); Green circle: Mannose.

The 2D docking profiles and associated non-covalent interactions were visualized by Discovery Studio, and the amino acid sites of molecular docking were obtained (Table 1). Venn diagram-based analysis was conducted on the intersection of the docking sites of the two ligands (Fig. 3). It was found that although no overlapping binding sites were observed between Asn-GlcNAc and Thr-Leu-Asn-GlcNAc in the molecular docking results, their respective binding sites were present in the cross-combination of other ligands (the amino acid residues in blue box). Following Venn diagram-based analysis, these amino acid residues were candidate sites: LYS-238, TRP-244, TRP-369, ARG-458, TRP-517, GLU-518, ARG-529, and THR-533. These amino acid residues were selected for single-amino-acid site-directed mutagenesis studies.

**Figure 3.**
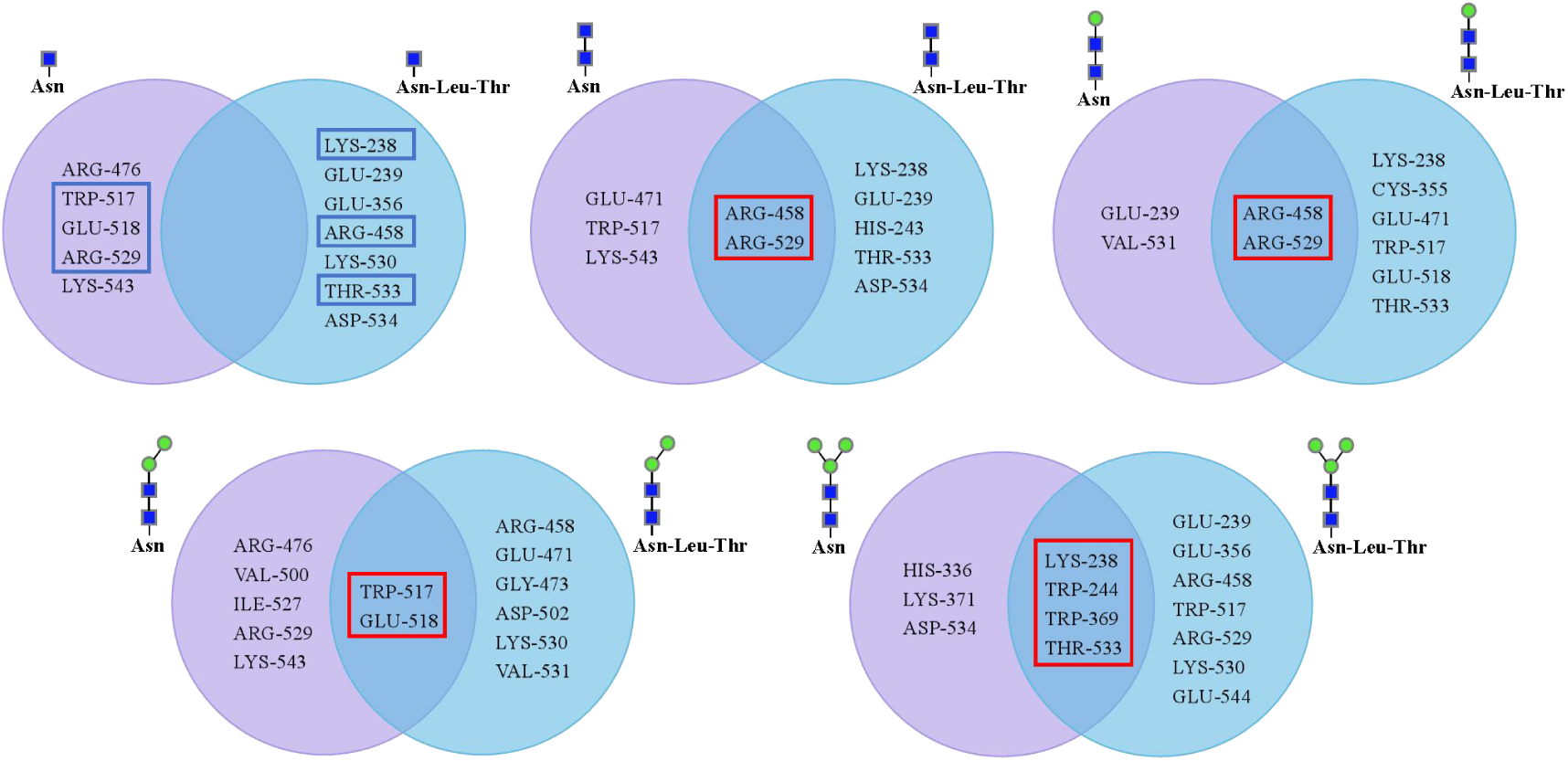
Venn diagram-based intersection screening of the docking amino acid residues. Asn and Asn-Leu-Thr were crossed by connecting two ligands of the same glycan chains respectively. The intersection amino acid residues were marked by red boxes, and the blue box showed the existence of the amino acid residues in the red boxes in the first Venn diagram.

**Table 1.**
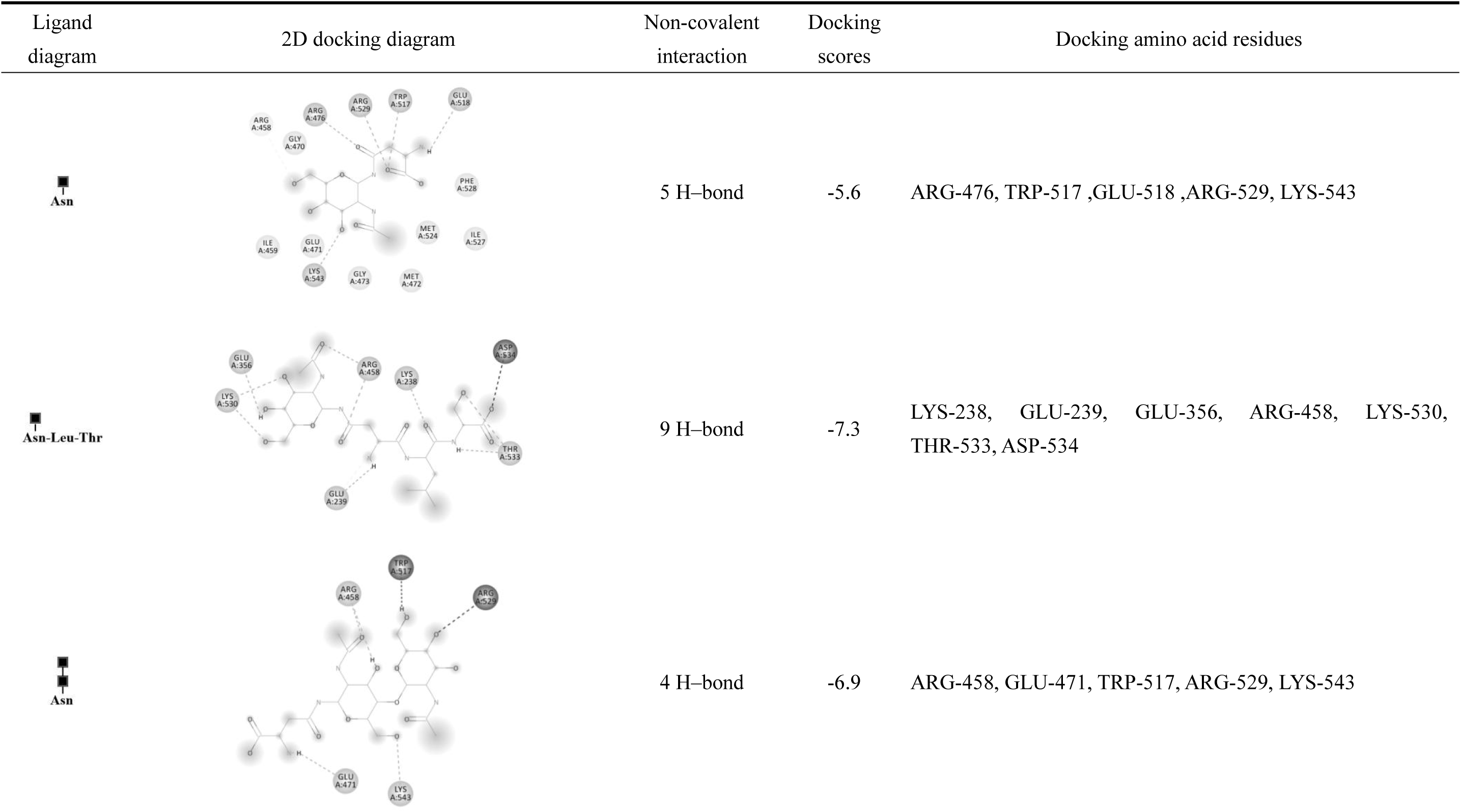
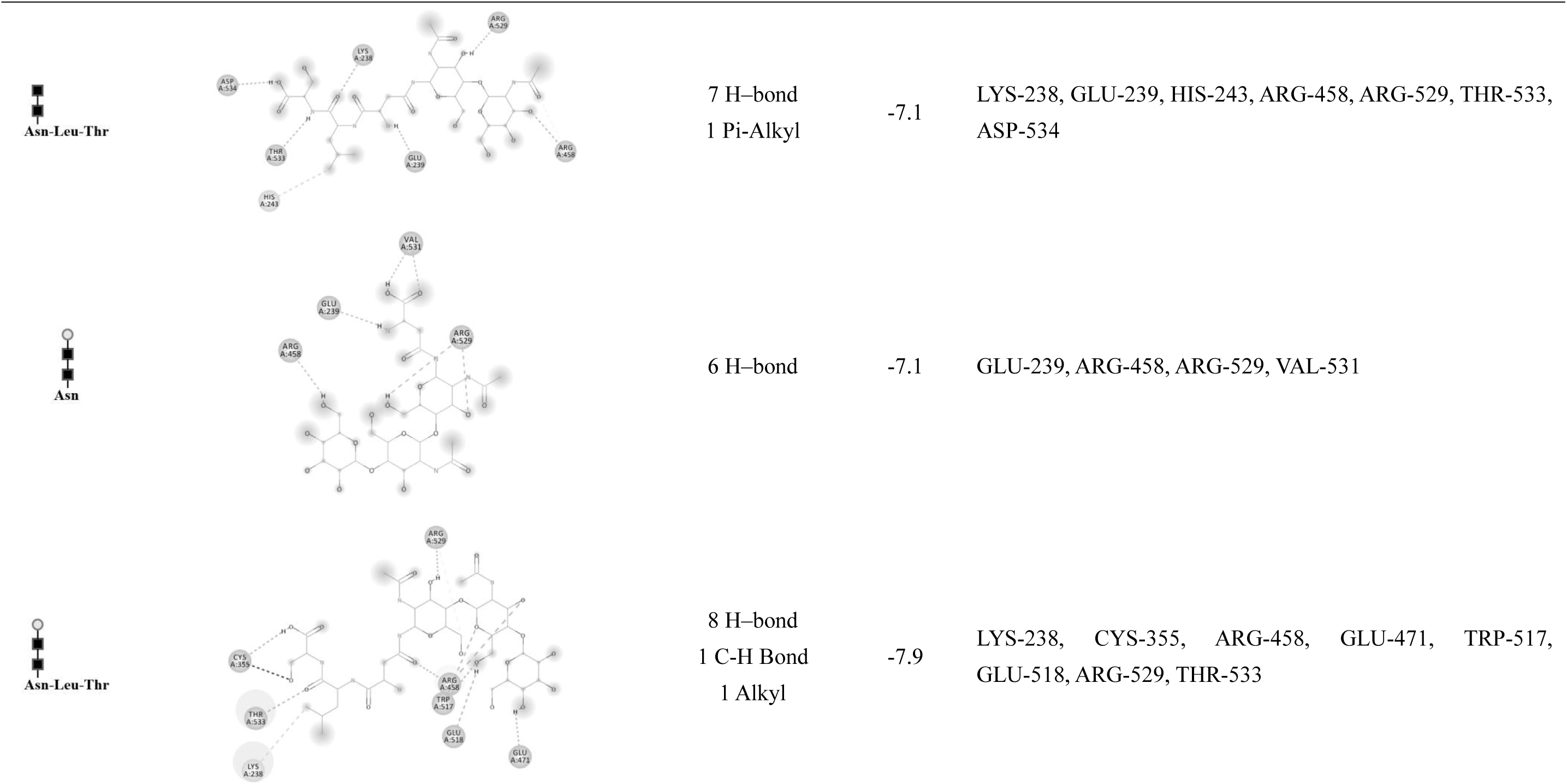
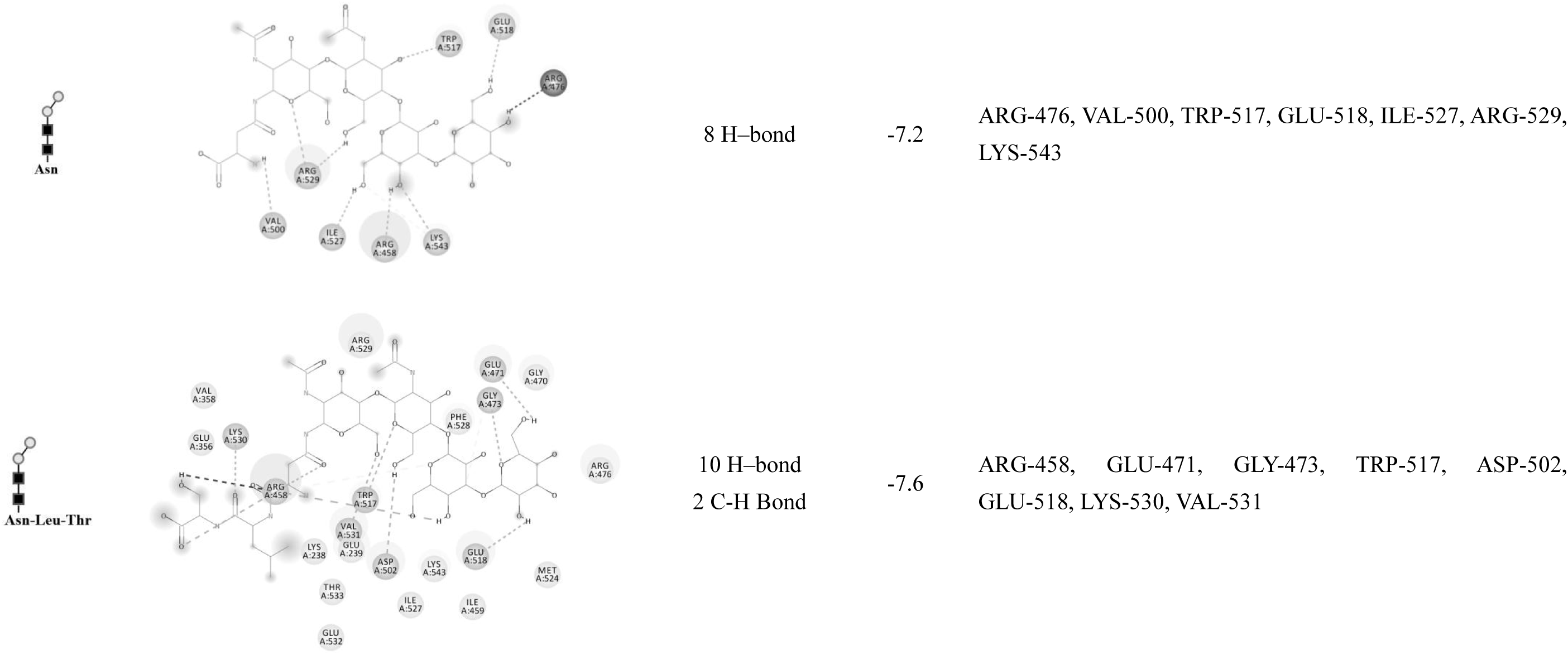
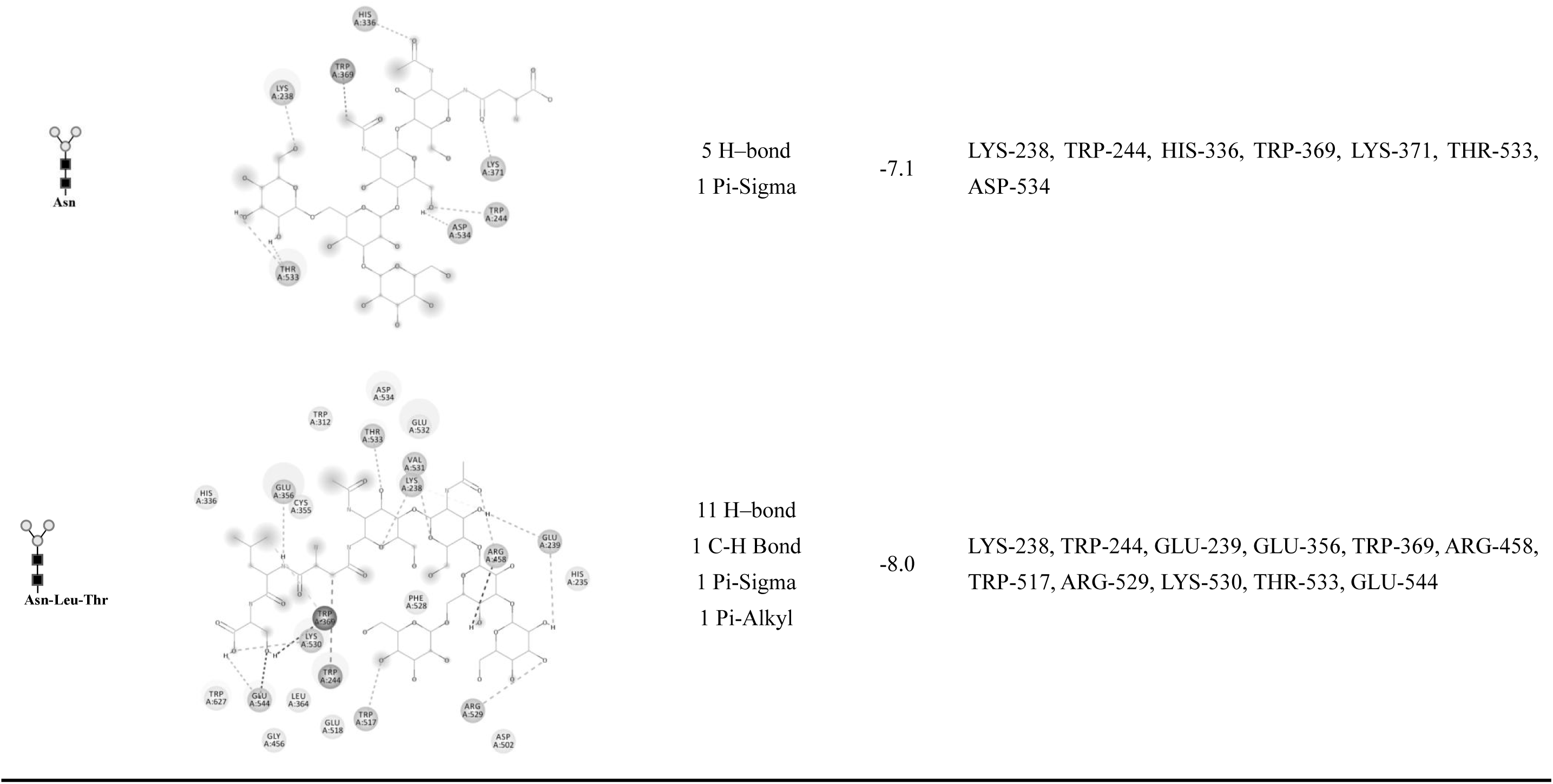
Sites of various ligand binding to NGLY1.

| Ligand diagram | 2D docking diagram | Non-covalent interaction | Docking scores | Docking amino acid residues |
| --- | --- | --- | --- | --- |
| Asn |  | 5 H-bond | -5.6 | ARG-476, TRP-517 ,GLU-518 ,ARG-529, LYS-543 |
| Asn-Leu-Thr |  | 9 H-bond | -7.3 | LYS-238, GLU-239, GLU-356, ARG-458, LYS-530, THR-533, ASP-534 |
| Asn |  | 4 H-bond | -6.9 | ARG-458, GLU-471, TRP-517, ARG-529, LYS-543 |

**Table1. (continued)**
| Ligand diagram | 2D docking diagram | Non-covalent interaction | Docking scores | Docking amino acid residues |
| --- | --- | --- | --- | --- |
| Asn-Leu-Thr |  | 7 H-bond<br>1 Pi-Alkyl | -7.1 | LYS-238, GLU-239, HIS-243, ARG-458, ARG-529, THR-533, ASP-534 |
| Asn |  | 6 H-bond | -7.1 | GLU-239, ARG-458, ARG-529, VAL-531 |
| Asn-Leu-Thr |  | 8 H-bond<br>1 C-H Bond<br>1 Alkyl | -7.9 | LYS-238, CYS-355, ARG-458, GLU-471, TRP-517, GLU-518, ARG-529, THR-533 |

**Table1. (continued)**
| Ligand<br>diagram | 2D docking diagram | Non-covalent<br>interaction | Docking<br>scores | Docking amino acid residues |
| --- | --- | --- | --- | --- |
| Asn |  | 8 H-bond | -7.2 | ARG-476, VAL-500, TRP-517, GLU-518, ILE-527, ARG-529, LYS-543 |
| Asn-Leu-Thr |  | 10 H-bond<br>2 C-H Bond | -7.6 | ARG-458, GLU-471, GLY-473, TRP-517, ASP-502, GLU-518, LYS-530, VAL-531 |

**Table1. (continued)**
| Ligand<br>diagram | 2D docking diagram | Non-covalent<br>interaction | Docking<br>scores | Docking amino acid residues |
| --- | --- | --- | --- | --- |
| Asn |  | 5 H-bond<br>1 Pi-Sigma | -7.1 | LYS-238, TRP-244, HIS-336, TRP-369, LYS-371, THR-533, ASP-534 |
| Asn-Leu-Thr |  | 11 H-bond<br>1 C-H Bond<br>1 Pi-Sigma<br>1 Pi-Alkyl | -8.0 | LYS-238, TRP-244, GLU-239, GLU-356, TRP-369, ARG-458, TRP-517, ARG-529, LYS-530, THR-533, GLU-544 |

**Table 2.**
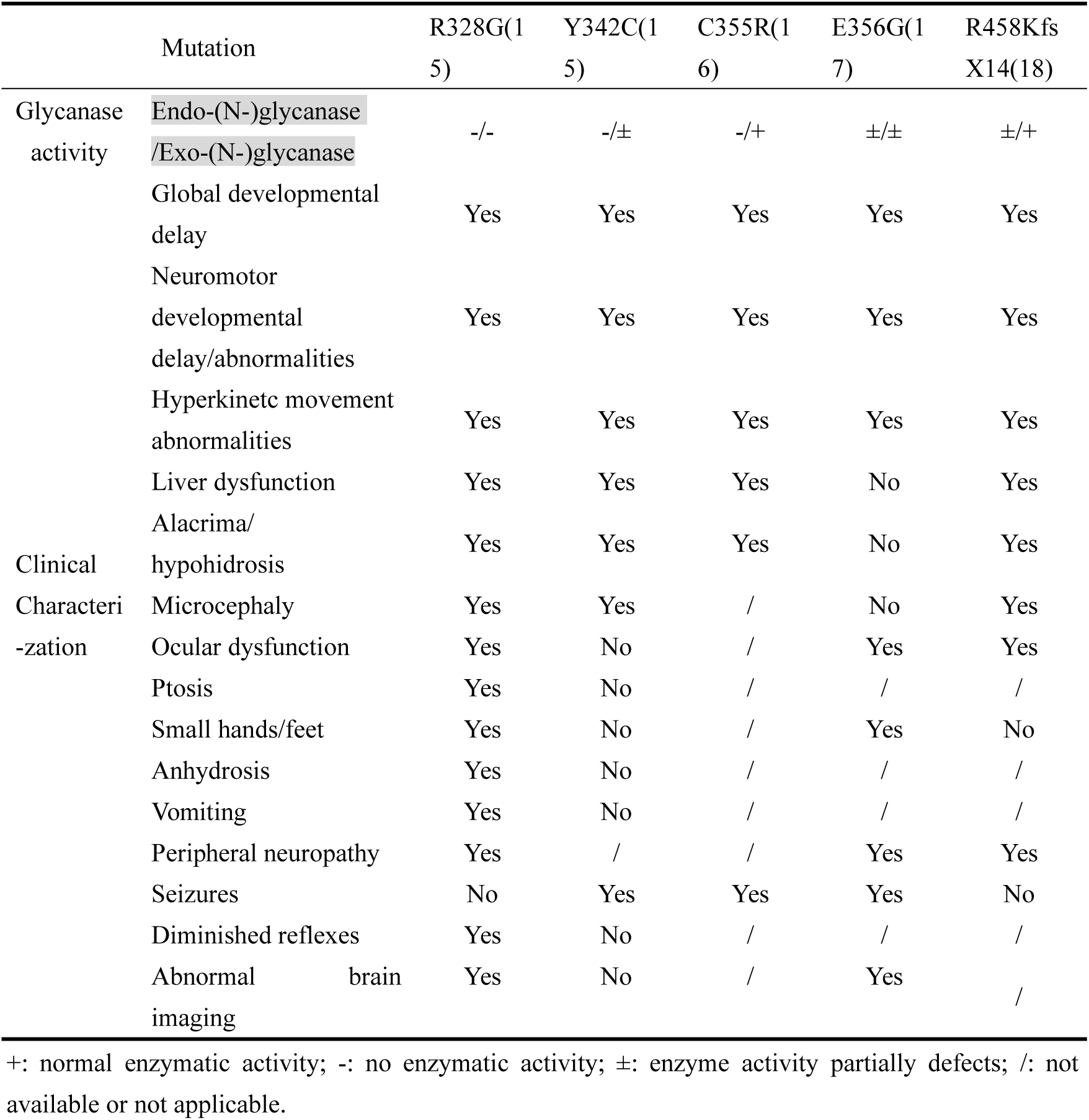
The activity of NGLY1 mutants and NGLY1-CDDG patients’ clinical characterization.

### The amino acid residues associated with exo-(N-)glycanase and endo-(N-)glycanase activity of NGLY1

Based on the 8 predicted active sites obtained through screening, the amino acids were specifically mutated to the amino acids with opposite charges. The successful construction of mutants was determined by sequencing of the recombinant plasmid genes (Fig. S1). The mutants of NGLY1 (K238D, W244R, W369E, R458E, W517R, E518K, R529E and T533A) were purified by Ni-NTA affinity chromatography.

The exo-(N-)glycanase activity of K238D and E518K were affected. Compared with the NGLY1-WT group, exo-(N-)glycanase activity of K238D was completely affected (Fig. 4A) and E518K exhibited lower exo-(N-)glycanase activity (Fig. 4B). Regarding endo-(N-)glycanase activity, K238D lacked endo-(N-)glycanase activity while E518K retained it.

**Figure 4.**
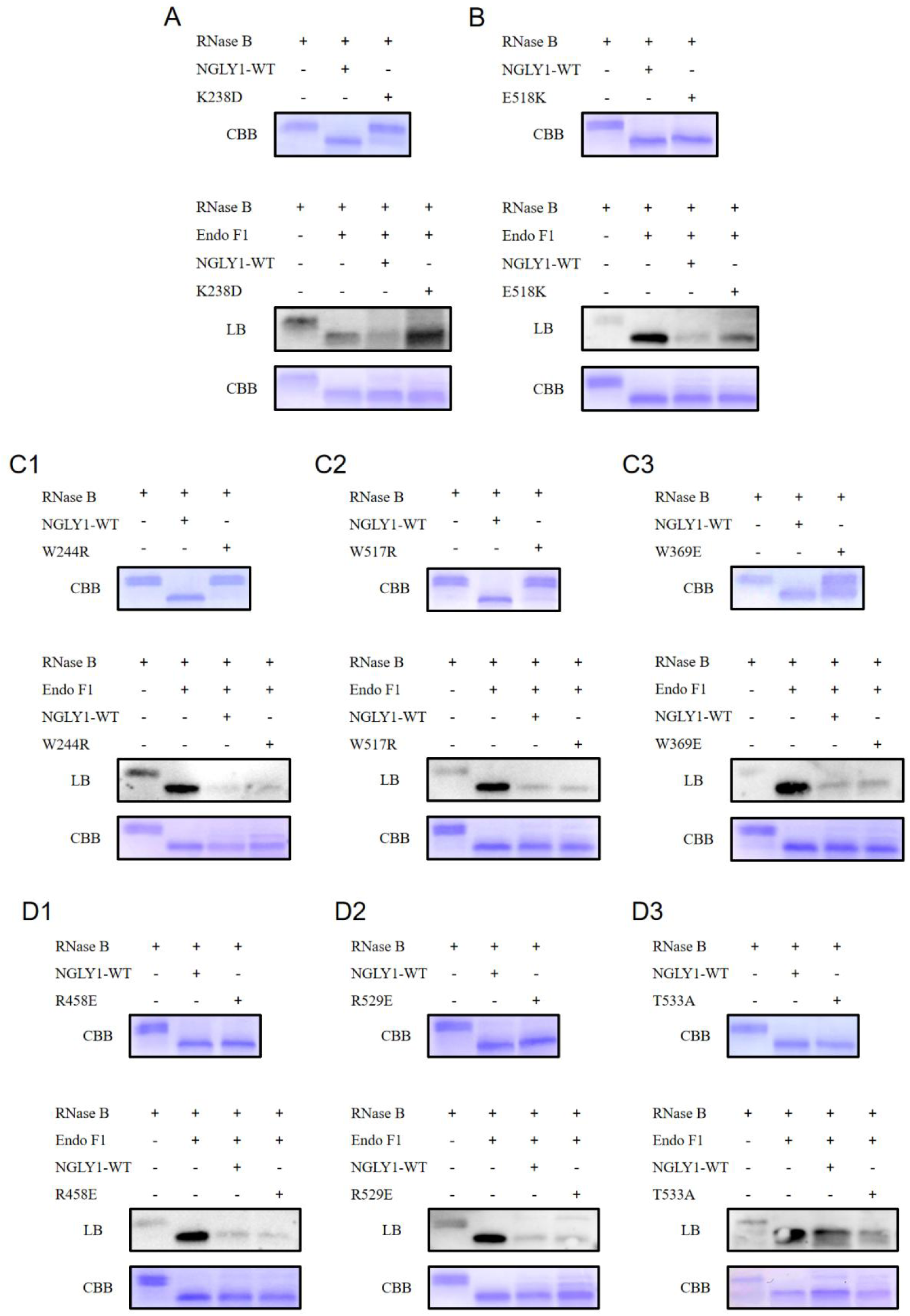
SDS-PAGE and lectin blot results of NGLY1 mutations’ endo-(N-)glycanase activity and exo-(N-)glycanase activity. A. K238D retains some endo-(N-)glycanase activity and has no exo-(N-)glycanase activity. B. E518K has endo-(N-)glycanase activity and retains some exo-(N-)glycanase activity. C1. W244R has no endo-(N-)glycanase activity and retains exo-(N-)glycanase activity. C2. W517R has no endo-(N-)glycanase activity and retains exo-(N-)glycanase activity. C3. W369E retains some endo-(N-)glycanase activity and has exo-(N-)glycanase activity. D1. R458E has endo-(N-)glycanase activity and exo-(N-)glycanase activity. D2. R529E has endo-(N-)glycanase activity and exo-(N-)glycanase activity. D3. T533A has endo-(N-)glycanase activity and exo-(N-)glycanase activity.

The exo-(N-)glycanase activity of W244R, W517R and W369E were not affected but their endo-(N-)glycanase activity were influenced (Fig. 4C1 and 3C2 and 3C3). The enzymatic activities of W244R and W517R towards RNase B were lost, whereas W369E had a lower enzymatic activity against RNase B.

In addition, it was found that the remaining screening intersection sites R458E, R529E and T533A had no effect on endo-(N-)glycanase activity nor exo-(N-)glycanase activity (Fig. 4D1, 4D2 and 4D3).

### The correlation between NGLY1’s exo-(N-)glycanase activity *in vitro* and NGLY1-CDDG’s clinical characterization

In this study, the effects of several patient mutants on the activities of NGLY1 endo-(N-)glycanase activity and exo-(N-)glycanase activity were investigated. The R328G mutant was completely inactive towards RNase B and the single GlcNAc, which meant R328G had no endo-(N-)glycanase activity nor exo-(N-)glycanase activity(Fig. 5A). The Y342C and E356G mutants had partial defects in the enzymatic activity of RNase B, and compared with the NGLY1-WT group, their exo-(N-)glycanase activity also had defects (Fig. 5B and 5D). The C355R mutant caused the loss of enzymatic activity of RNase B, while maintaining enzymatic activity towards a single GlcNAc (Fig. 5C).

**Figure 5.**
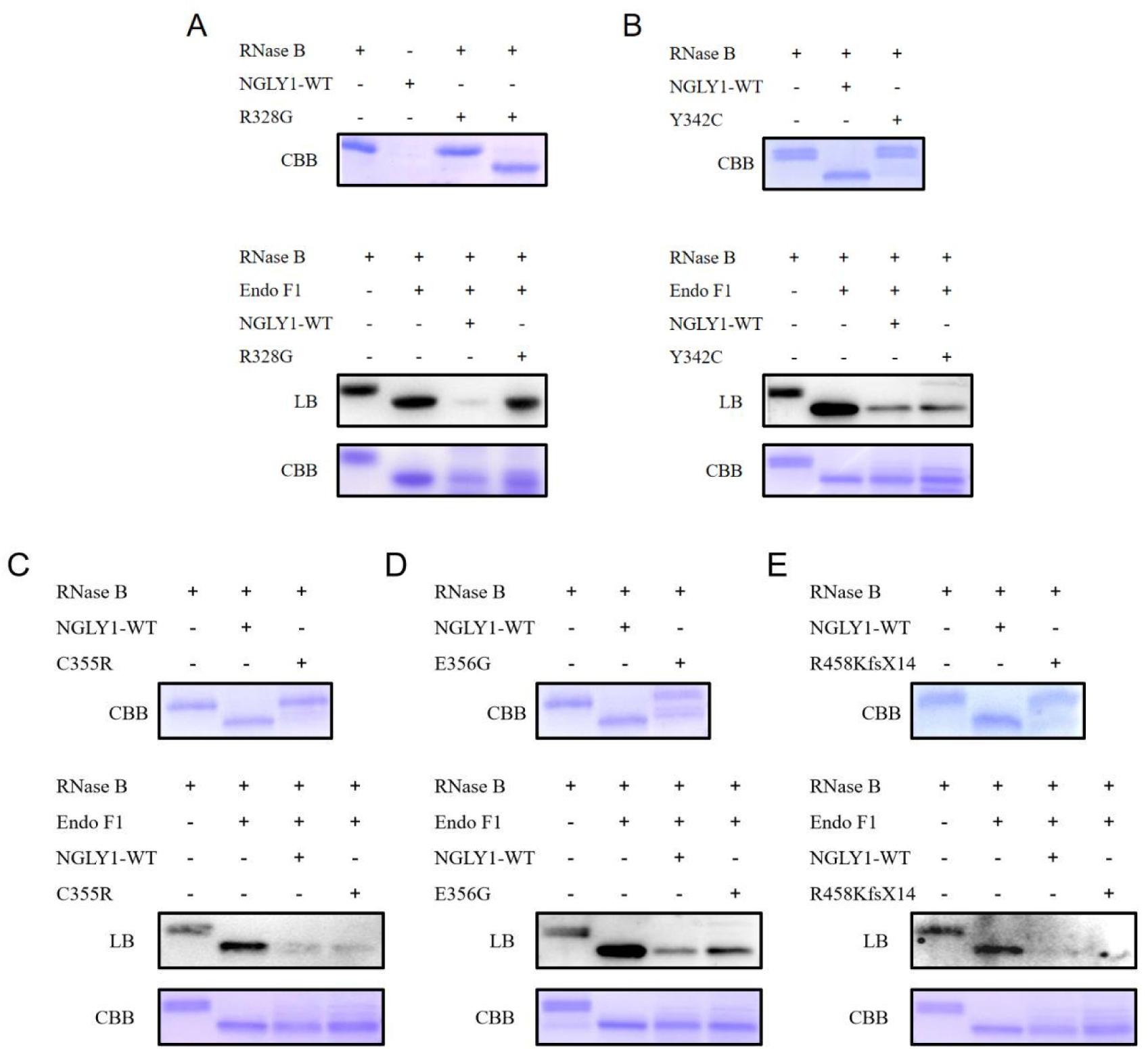
The endo-(N-)glycanase activity and exo-(N-)glycanase activity of the reported NGLY1 patients’ mutations. A. R328G has no endo-(N-)glycanase activity nor exo-(N-)glycanase activity. B. Y342C remains some endo-(N-)glycanase activity and has no exo-(N-)glycanase activity. C. E356G remains some endo-(N-)glycanase activity and has no exo-(N-)glycanase activity. D. C355R have no endo-(N-)glycanase activity and have exo-(N-)glycanase activity. E. R458KfsX14 remains some endo-(N-)glycanase activity and has exo-(N-)glycanase activity.

Furthermore, the clinical symptoms of these NGLY1-CDDG patients and the endo-(N-)glycanase activity and exo-(N-)glycanase activity of their mutants were summarized (Table2). The exo-(N-)glycanase and endo-(N-)glycanase activities of R328G were completely absent. Correspondingly, the symptoms of patients with the R328G mutation appeared to be more severe, with additional symptoms such as abnormal brain imaging, diminished reflexes, anhidrosis, vomiting, ocular dysfunction, ptosis and small hands/feet compared to Y342C patients. This might be because, the NGLY1 in Y342C mutation patients, although mutated, still retained some exo-(N-)glycanase activity, which could remove N-GlcNAc proteins in body to a certain extent, thereby exerting a detoxification effect. The exo-(N-)glycanase activity of C355R was completely retained, and the symptoms of patients with the C355R mutation were slightly milder. For instance, symptoms such as microcephaly, ocular dysfunction, abnormal brain imaging, and anhidrosis have not been reported in C355R patients. E356G not only retained partial exo-(N-)glycanase activity but also partial endo-(N-)glycanase activity. Its symptoms were milder than those of other patients, with no liver dysfunction, alacrima/hypohidrosis, or microcephaly. R458KfsX14 retains some endo-(N-)glycanase activity and completely retains exo-(N-)glycanase activity. The patients with this genotype did not exhibit symptoms of small hands/feet or seizures. It was speculated that the dual glycanase activity of NGLY1 might be functioning simultaneously in the patients’ body.

## Discussion

Previously, it has been proposed that NGLY1 is a key component of the ERAD process for newly synthesized proteins. This process was designated as ER-assisted digestion of improperly folded glycoproteins(19). By predicting protein interactions through the STRING website (https://cn.string-db.org/), a connection between NGLY1 and ENGase was found (Fig. S3). *Ngly1* knockout studies conducted in mice demonstrated that the embryonic lethal phenotype of *Ngly1* could be rescued by a second knockout of *Engase*(10). Previous studies have demonstrated that upon compromised Ngly1 activity, ENGase-mediated deglycosylation of misfolded glycoproteins may cause excess formation of N-GlcNAc proteins in the cytosol(11). Together, these studies and our present work indicate that since both ENGase and NGLY1 work on the same or similar substrates, the down-regulation of NGLY1 may stimulate an up-regulation of ENGase and generate more ENGase products (N-GlcNAc proteins), releasing the ERAD stress caused by NGLY1 deficiency. We hypothesized that this activity could partially compensate for the deglycosylation of misfolded glycoproteins mediated by ENGase. The accumulation of N-GlcNAc proteins could result in the aggregation and elimination of functional proteins within the cell. Based on early studies and our own results generated from *C. elegans*, we hypothesized that ENGase products may represent an index of stress, including the ones from environmental and genetics(20). Therefore, we suggested that there may be an exo-(N-)glycanase activity that could relieve the protein stress resulting from increased ENGase activity.

An *in vitro* assay was conducted for this activity by using the ENGase products as substrates. To our surprise, an exo-(N-)glycanase activity of NGLY1 was detected. In addition, the similar activity was also detected in PNGase F and PNGase F-Ⅱ from *E. meningoseptica.* This study provided the first confirmation at the level of proteins that PNGase F, PNGase FII and NGLY1 can effectively hydrolyze N-GlcNAc proteins. The study by Fan JQ and Lee YC (1997) demonstrated that PNGase F exhibited weak enzymatic activity against two synthetic GlcNAc-peptides (a tripeptide and a pentapeptide)(21). In comparison with that previous report of PNGase F’s hydrolytic activity toward N-GlcNAc peptides, this study significantly expands the range of intact protein substrates for NGLY1’s exo-(N-)glycanase activity. Not only does it reveal broader biochemical functions, it also provides new research dimensions and a critical theoretical foundation for deepening our understanding of biological significance of NGLY1. We also illustrated that NGLY1, PNGase F and PNGase F-II had no enzymatic cleavage effect on single amino acid linked with GlcNAc (Asn-GlcNAc) (Fig. S2), suggesting that peptide N-glycanase exerted exo-(N-)glycanase activity by recognizing N-GlcNAc on proteins only. In addition, time-gradient experiments were conducted for the two enzymatic activities of NGLY1, respectively. The results revealed a time-dependent enhancement of its exo-(N-)glycanase activity with increasing digestion duration (Fig. S4). Hirayama established a quantitative, non-radioisotope (RI)-based assay method for measuring recombinant NGLY1 using a BODIPY-labeled asialoglycopeptide (BODIPY-ASGP) derived from hen eggs(22), representing a significant methodological advance. Our study revealed that NGLY1 exhibited both established endo-(N-)glycanase activity and a previously unrecognized exo-(N-)glycanase function, and future investigations will explore the applicability of these reported quantitative methods to NGLY1’s exo-(N-)glycanase activity.

In this study, the discovery of NGLY1’s exo-(N-)glycanase activity has created a new double-hit mechanism for NGLY1-CDDG. *In vitro* results generated from NGLY1-CDDG associated mutations were consistent with the dual-function model. The defects in endo/exo-glycanase of NGLY1 were detected in most of the mutations, indicating a correlation between NGLY1’s exo-(N-)glycanase activity and clinical symptoms of NGLY1-CDDG.

In a previous study, the active sites of NGLY1 were predicted, and the mutations on these sites were listed as high risk mutants associated with rare disease NGLY1-CDDG(23). In this study, additional high-risk sites were added based on the results generated from assays for exo-(N-)glycanase activity of NGLY1. The addition may provide a new reference value for the diagnosis and treatment of NGLY1-CDDG. Firstly, the results generated from the dual enzyme activities assay established in this study could be used for subtyping of NGLY1-CDDG. Secondly, although our data demonstrated that NGLY1-CDDG were associated with defects of exo-(N-)glycanase activity of NGlY1, the results did not exclude the possibility that other enzymes may also involve in the detoxification process for N-GlcNAc proteins. The possibilities are under the investigation. Quantitative analysis of both enzymatic activities is of great significance for NGLY1-CDDG’s characterization and provides novel evidence to advance research and precision diagnostics for NGLY1-related disorders, although further methodological development is warranted. Finally, the predicted active site mutations without reported NGLY1-CDDG may open a discussion on other symptoms or diseases associated with this gene.

### Experimental procedures

#### Purification of NGLY1

The NGLY1 protein inducible expression and purification system has been established and the methods used in this study were based on the previously published articles of our research group(23,24).

#### Detection on endo-(N-)glycanase activity of NGLY1

The enzyme activity assay of NGLY1 has been established in advance(23). NGLY1 (2 μg) was mixed with RNase B (New England Biolabs, Ipswich, Massachusetts, USA) (2 μg) previously denatured at 100 ° C, and add PBS to make the total reaction system (20 μL). Incubate at 37° C for 12-16 hours, then separate by 15% SDS-PAGE and conduct Coomassie blue staining. The molecular weight of RNase B is approximately 17 kDa. After NGLY1-mediated de-sulfation, the molecular weight of RNase B drops to approximately 13.7 kDa. When the activity of NGLY1 is inhibited, the molecular weight of RNase B remains at approximately 17 kDa.

#### Detection on exo-(N-)glycanase activity of NGLY1

The product of RNase B cleaved by Endo F1 was used as the substrate for verifying the NGLY1’s exo-(N-)glycanase activity. After RNase B (2 μg) was subjected to a 100 ° C denaturation treatment for 10 minutes, Endo F1 (2 μg) was added for enzymatic digestion for 12-16 hours at 37° C. After 10 minutes of denaturation at 100 ° C, it was cooled on ice. Equal amounts of enzyme (4 μg) were added, and enzymatic digestion is carried out for 12-16 hours.

Then, loading buffer was added for denaturation treatment.

Through the SDS-PAGE electrophoresis method, the proteins in the sample were separated according to their molecular weights. The separated proteins were transferred onto a PVDF membrane. The transfer conditions were 220V for 100 minutes. The membrane was treated with 5% bovine serum albumin (Beyotime, Shanghai, China) blocking solution at room temperature on a shaker for 2 hours to block non-specific binding sites. The membrane was incubated with 1:200 diluted Wheat Germ Agglutinin (WGA) (Vector Laboratories, San Francisco Bay Area, USA) at 4° C on a shaker overnight (12-16 hours) to allow the lectin to bind to the target carbohydrate chains on the membrane proteins. After the lectin fully binds, the membrane was washed 1x TBST for 10 minutes, repeated three times, and the free lectin is removed. Then, 1:2000 diluted horseradish peroxidase-labeled Streptavidin (Beyotime, Shanghai, China) was added at room temperature on a shaker for 2 hours. 1× TBST is washed for 10 minutes, repeated three times, and the ECL chemiluminescent substrate reagent (biosharp, Hefei, Anhui, China) was added to the membrane. The fluorescence signal was detected by luminescence imaging to determine the presence and relative content of the target carbohydrate chain.

#### Mass spectrometry analysis on exo-(N-)glycanase activity of NGLY1

Mass spectrometry analysis was conducted using the glycopeptide EEQYN(GlcNAc)STYR (a gift from Professor Xinmiao Liang’s research group, Dalian Institute of Chemical Physics, Chinese Academy of Sciences). A 20 μL reaction mixture containing the glycopeptide, NGLY1, and buffer was incubated at 37 °C for 48 h. The resulting mixture was desalted, lyophilized, and redissolved in 50% ACN. After mixing with DHB matrix solution, samples were analyzed on a RapifleX MALDI-TOF mass spectrometer (Bruker) equipped with a 337 nm laser. Spectra were acquired in the m/z range of 650-3200 Da with a resolution of 20,000, accumulating 2000 laser shots.

#### Computer virtual molecular docking of NGLY1 and ligands

The three-dimensional structure of NGLY1 was previously predicted using the amino acid sequence of NGLY1 through the online tool AlphaFold2 (website: https://colab.research.google.com/github/sokrypton/ColabFold/blob/main/AlphaFold2.ipynb?pli=1#scrollTo=kOblAo-xetgx, accessed on January 10, 2022).

The three-dimensional structures of 10 ligands were drawn, generated the 3D structures in OpenBabel, and removed water molecules and hydrogen atoms. Using the "AutoDock vina" function in AutoDock software, we performed molecular docking of these 10 ligands with NGLY1 respectively. NGLY1 consists of three domains: the PUB domain, the PNG core domain, and the PAW domain. Among them, the PNG core domain plays the main catalytic role. Center the receptor pocket around the PNG core domain of the NGLY1 protein (coordinates x = -4.054, y = -4.87, z = 2.613), and set this receptor pocket as a cubic grid with side lengths (113×70×112A). AutoDock Vina uses BFGS local optimization and genetic algorithms to generate 9 conformations. The lowest energy conformation was selected and their interaction with NGLY1 were analyzed using the PyMOL (version 3.1.3).

#### Construction of NGLY1 amino acid mutation and enzymatic activity analysis

The pET28a-NGLY1 (wild type) plasmid is kept in our laboratory. All NGLY1 mutants were constructed using the pET28a-NGLY1 (wild type) plasmid as the template and reverse complementary primers through PCR (Takara, Beijing, China). The constructed pET28a-NGLY1 (wild type) plasmid and pET28a-NGLY1 (mutant) plasmids were respectively transformed into Escherichia coli BL21(DE3) (Tiangen, Beijing, China) for protein expression. The primer sequences are shown in Table 3. The purification of NGLY1 mutants and the detection methods of endo-(N-)glycanase and exo-(N-)glycanase activities were the same as those for NGLY1.

**Table 3.**
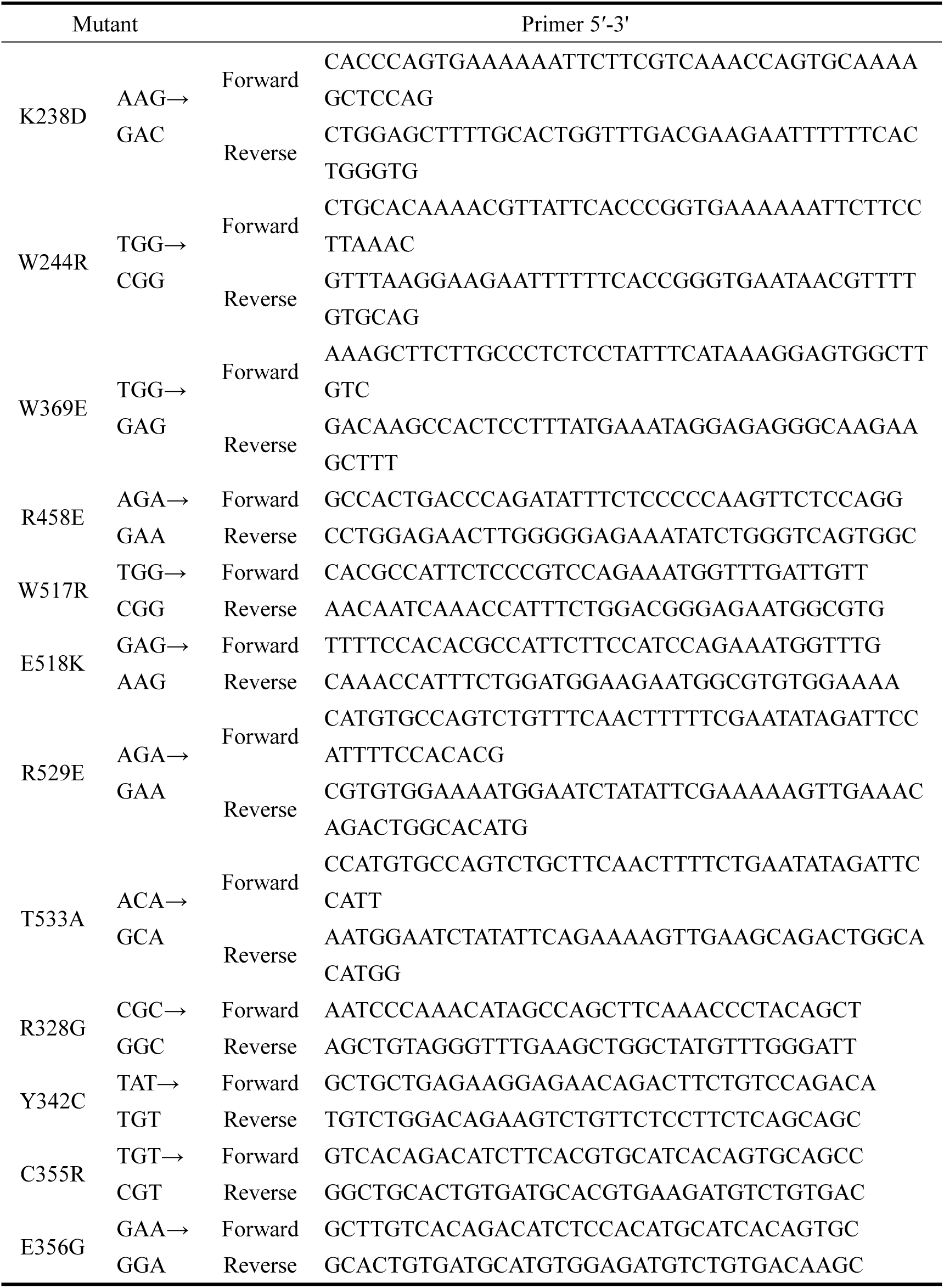
PCR primer sequences for constructing NGLY1 mutants.

## Data availability

All data are included in the manuscript and supporting information.

## Supporting Information

This article contains supporting information.

## Funding and additional information

This work was supported by Open Research Project of the National Key Laboratory for Genetic Regulation of Complex Traits (SKLGE-2314).

We honestly thank Xinmiao Liang, Cuiyan Cao (Dalian Institute of Chemical Physics, Chinese Academy of Sciences) and Liming Wei (Institutes of Biomedical Sciences, Fudan University) for suggestions.

