## Supporting Information for "Identification and characterization of a novel exo-(N-)glycanase activity of NGLY1 on ENGase-digested N-GlcNAc proteins *in vitro*"

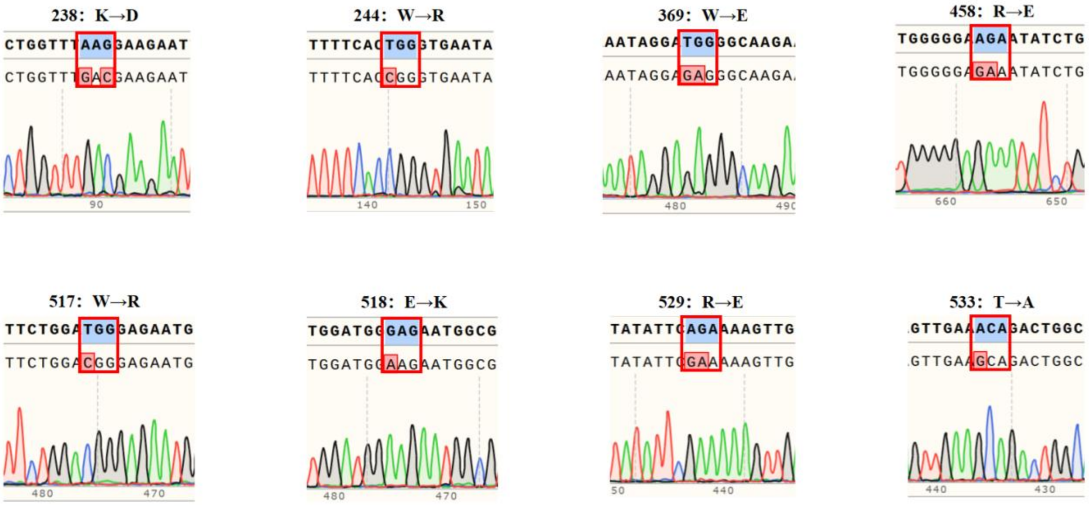


**Supplementary Figure 1. Sequence of NGLY1 mutants' recombinant plasmid genes.** The numbers indicated the amino acid sites of the mutations, and the mutated bases were marked with red boxes.

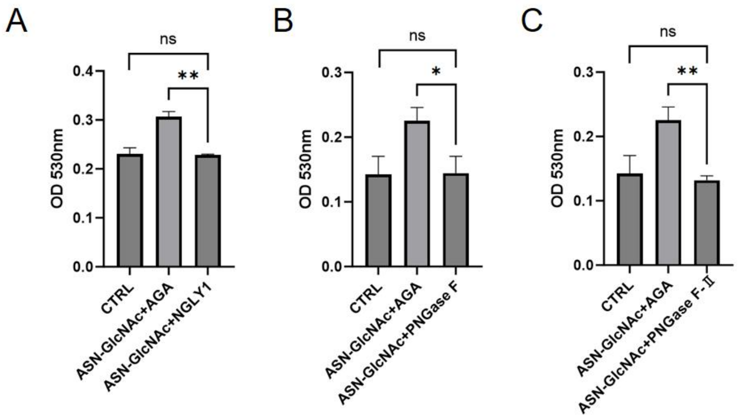


**Supplementary Figure 2. Analysis of NGLY1's enzymatic cleavage effect on Asn-GlcNAc.** A. NGLY1’s enzymatic activity effects on Asn-GlcNAc. B. PNGase F’s enzymatic activity effects on Asn-GlcNAc. C. PNGase F-II’s enzymatic activity effects on Asn-GlcNAc. CTRL: control; AGA: aspartyl N-acetylglycylaminoglucomannosidase (positive control enzyme); ns: no significant; *:$P < 0.05$; **: $P < 0.01$. Aspartyl N-acetylglycylaminoglucomannosidase（AGA）could hydrolyze GlcNAc-Asn , generating a single GlcNAc and aspartic acid (Asp)(1).

1. Kaartinen, V., Mononen, T., Laatikainen, R., and Mononen, I. (1992) Substrate specificity and reaction mechanism of human glycoasparaginase. The N-glycosidic linkage of various glycoasparagines is cleaved through a reaction mechanism similar to L-asparaginase. *J Biol Chem* **267**, 6855-6858

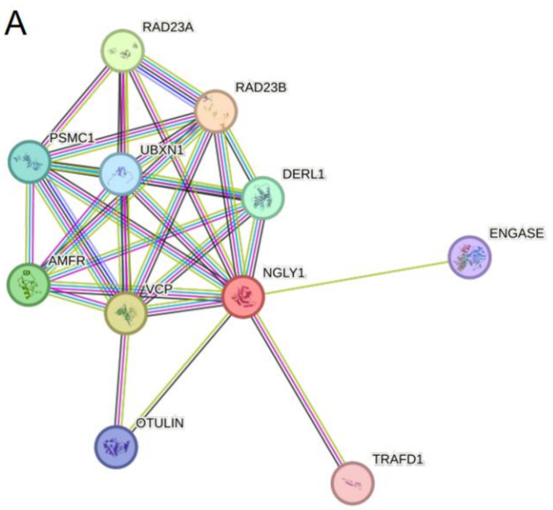


**Supplementary Figure 3. Interaction network of NGLY1.** Edges represent protein-protein associations. Associations are meant to be specific and meaningful, proteins jointly contribute to a shared function; this does not necessarily mean they are physically binding to each other. A. A connection between NGLY1 and ENGase was found from textmining.

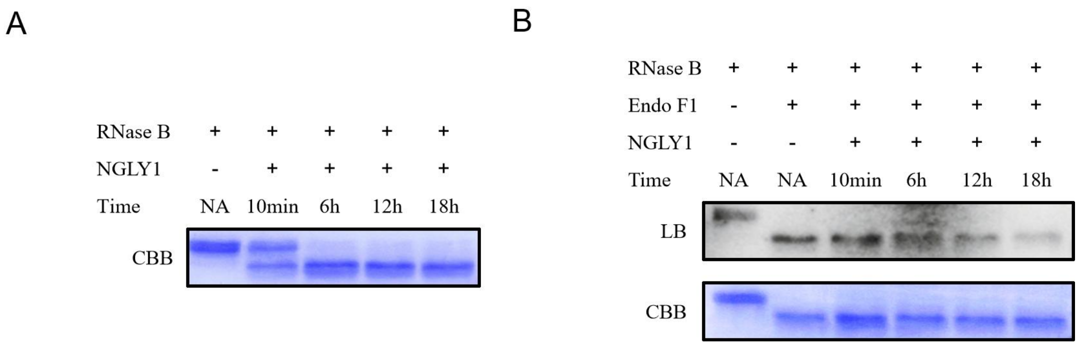


**Supplementary Figure 4. Time-course dual-enzyme digestion validation of NGLY1.** Time-course experiments for both endo-(N-)glycanase and exo-(N-)glycanase activities have been performed at time points of 10 min, 6 h, 12 h, and 18 h, with samples collected at each interval and analyzed together by SDS-PAGE and lectin blot. A. Time-course validation of NGLY1 endo-(N-)glycanase activity. B. Time-course validation of NGLY1 exo-(N-)glycanase activity. NA: Not Applicable.
